# “Plasmids drive the dissemination and diversification of Type VI Secretion Systems”

**DOI:** 10.64898/2025.12.03.692189

**Authors:** María del Mar Quiñonero-Coronel, M. Pilar Garcillán-Barcia

**Affiliations:** Instituto de Biomedicina y Biotecnología de Cantabria (IBBTEC), Consejo Superior de Investigaciones Científicas-Universidad de Cantabria, Santander, Spain

**Keywords:** Type VI secretion system, plasmids, horizontal gene transfer

## Abstract

The Type VI secretion system (T6SS) is a major determinant of bacterial competition, yet its dissemination across lineages remains unclear. Analyzing 43,213 plasmids and 29,161 chromosomes, we reveal plasmids as an underestimated reservoir and vehicle for T6SS diversification. We identified 405 complete plasmidencoded T6SSs and 929 orphan islands containing *hcp*, *vgrG*, and/or *PAAR* genes, often independent of full systems. Plasmid-encoded T6SSs are biased toward large replicons, frequently megaplasmids, with distinct stability and mobility traits: orphan island plasmids are enriched in conjugation modules, whereas complete systems rely on partition and toxin–antitoxin maintenance. Phylogenomic analyses show some plasmid lineages stably integrating T6SSs as core traits, while others undergo recurrent acquisition and diversification. Comparative and ancestral analyses indicate pervasive bidirectional transfers between plasmids and chromosomes, mediated by insertion sequences. The presence of near-identical homologs across compartments underscores the capacity of plasmids to transcend phylogenetic barriers and propagate these nanoweapons. Together, our results identify plasmids as dual evolutionary actors in T6SS ecology—short-term vectors that enable rapid horizontal spread, and long-term reservoirs that foster stabilization and adaptive diversification.

## Introduction

Bacteria have evolved a remarkable array of secretion systems to interact with their environment, ranging from establishing symbiosis to mediating interbacterial competition [1]. Among these, T6SS has emerged as a widespread and versatile nanomachine, capable of delivering toxic effectors into neighboring cells and modulating microbial community structure [2–4]. T6SSs contribute to both antagonistic and cooperative interactions, influencing bacterial survival, adaptation, and ecological fitness [5, 6].

Traditionally, T6SSs are found to be encoded in chromosomes of Gram-negative bacteria [7], where their evolutionary history and functional roles have been extensively studied. However, T6SSs can also be found on plasmids [8], although their functionality has been scarcely tested [9–12]. Because plasmids are inherently mobile and often cross species and genus boundaries [13], plasmid-encoded T6SSs represent potential vectors for the ecological redistribution of bacterial antagonism and cooperation. Their occurrence raises key questions about how mobile genetic elements (MGEs) shape interbacterial competition, niche colonization, and the coevolution of antagonistic systems with plasmid mobility.

Plasmids are central drivers of bacterial evolution, serving as reservoirs of adaptive traits such as antibiotic resistance, metabolic pathways, and virulence factors [14]. Through horizontal gene transfer they provide a rapid means for bacteria to acquire competitive functions [15]. Beyond their role as genetic vehicles, plasmids shape the tempo and mode of bacterial evolution by linking gene mobility with ecological opportunity. The acquisition of T6SSs by plasmids could broaden their ecological roles, promoting the spread of competitive capabilities across bacterial lineages [16]. However, the evolutionary origins, host range, and functional associations of plasmid-borne T6SSs remain poorly understood, as does whether their mobility results in transient acquisitions or enduring ecological strategies. From an ecological and evolutionary perspective, understanding how plasmids mediate the mobilization and stabilization of T6SSs is essential to explain how microbial populations balance horizontal gene flow with competitive stability.

To address these questions, we performed a comprehensive analysis of T6SSs across plasmids and chromosomes from diverse bacterial families. Using integrated network and phylogenomic approaches, we examined their distribution, evolutionary relationships, and associated functional traits. Our results reveal that distinct plasmid lineages specialize in the maintenance and diversification of T6SSs, positioning plasmids not merely as vehicles shuttling systems between genomes but as persistent reservoirs that co-evolve with specific bacterial hosts while occasionally bridging taxonomic boundaries. This lineage-level resolution highlights plasmids as short-term vectors that disseminate T6SS modules across taxa, but also as long-term reservoirs that maintain and diversify these systems within specific lineages, ultimately influencing community composition and ecosystem function.

## Materials and Methods

### Genome dataset

Complete genomes were retrieved from the NCBI RefSeq212 database (June 2022), including 29,161 chromosomes and 43,872 plasmids (Supplementary Tables S1–S2). To minimize misclassified entries, plasmids were curated following [17], yielding 43,213 plasmid sequences after filtering.

### T6SS detection and classification

T6SS clusters were identified in chromosomal and plasmid datasets using MacSyFinder v1.0.5 [18] with HMM profiles for subtypes i–iii. Detection parameters required ≥2 distinct T6SS genes within ≤20 intervening genes (--min-genes-required 2; --min-mandatory-genes-required 2; --inter-gene-max-space 20). Clusters with ≥8 components were classified as complete T6SSs and assigned to subtypes via diagnostic markers; others were labeled incomplete or as putative orphan islands if containing *tssD–tssI*, *tssD–evpJ*, *tssI–evpJ*, or *tssD–tssI–evpJ* gene pairs. Isolated TssI (also named VgrG) and TssD (also known as Hcp) proteins were detected using HMMERv3 (hmmsearch; --incE 1e-3, --incdomE 1e-3, ≥50% coverage of profile length). T6SS gene synteny was obtained with Clinker v0.0.31 [19].

### Plasmid classification

Plasmids were assigned to Plasmid Taxonomic Units (PTUs) with COPLA [20]. As previously defined by [21], the threshold for megaplasmid classification was set at ≥5% of the median genome size of the host taxonomic family (Supplementary Table S3), obtained from EZBioCloud [22].

### Plasmid functional annotation

Relaxases and conjugation systems were detected using MOBscan [23] and CONJscan [24]. Antimicrobial resistance (AMR) genes were identified with Staramr [25], using default parameters. Virulence factors were screened with ABRicate against VFDB [26] (https://github.com/tseemann/abricate; BLASTp, e-value <1e-5, ≥50% identity, ≥60% coverage). Insertion sequences were annotated via ISFinder [27]. Recombinases (tyrosine, serine, and resolvase domains; PF00589, PF07058, PF00239) were detected using HMMERv3 (hmmscan; --E 1e-5, --domE 1e-5, --incE 1e-5, --incdomE 1e-5, ≥50% coverage of profile length). Partition systems were searched with hmmsearch (-E 1e-5, --domE 1e-5, --incE 1e-5, --incdomE 1e-5; ≥50% profile coverage) using the HMM profiles for type I (PF01656, PF13614, PF18607, NF010259, PF08775, PF02195, PF07506, PF18090, PF08535, PF06613, PF09274), type II (PF06406, PF21523), and type III (PF21493) class components. HMM profiles for the toxins of the toxin-antitoxin (TA) systems contained in TADB3.0 [28] were retrieved using HMMERv3 (hmmscan; --E 1e-5, --domE 1e-5, --incE 1e-5, --incdomE 1e-5, ≥50% sequence coverage). These profiles (Supplementary Table S4) were then used to screen for putative TA modules in the plasmid dataset (hmmsearch; --E 1e-5, --domE 1e-5, --incE 1e-5, --incdomE 1e-5, ≥80% sequence coverage).

### Phylogenetic analyses

TssC proteins from clusters with ≥10 components were grouped with MMseqs2 15.6f452 [29] at 99% identity and 100% coverage, yielding 2,101 representatives. The subtype iv sheath (WP_012473177.1) from *Amoebophilus asiaticus* 5a2 was included. Alignments were generated with MAFFT v7.271 (--retree 2 --maxiterate 1000) [30], trimmed with TrimAl v1.2 (-automated1) [31] and used to construct a maximum-likelihood (ML) phylogeny in IQ-TREE [32] under the LG+F+R10 model [33], with 1,000 ultrafast bootstraps [34]. The tree was rooted with the T4 gp18 sheath subunit. Plasmid diversity within PTUs was assessed using kSNP v3.0 [35] with Kchooser-defined k-mers. Trees were reconstructed by maximum parsimony (-core option). All trees were visualized with iTOL v5 [36].

The inference of ancestral states for TssC genomic contexts (chromosomal or plasmid) was carried out with PastML v1.9.33 [37] using ML with the MPPA algorithm under the F81 model.

### Cumulative distribution function (CDFs)

Patristic distances were computed from the TssC tree using the cophenetic.phylo function in R (ape v5.6-2). For each plasmid-encoded TssC, the closest homolog (plasmid or chromosome) was identified. For clustered but contextually distinct sequences, a distance of zero was assigned. CDFs of patristic distances and plasmid sizes were generated and visualized with seaborn.ecdfplot (v0.11.0, Python v3.13). Statistics were performed using the one-sided Mann-Whitney U test calculated with scipy.stats.mannwhitneyu (v1.14.1, Python v3.13).

### Network construction

Monopartite networks of T6SS-encoding plasmid similarity were obtained for pairwise average nucleotide identity values with a 50% plasmid length threshold (ANI_L50_), calculated as described by [13]. Bipartite networks of chromosome–plasmid T6SS transfers were built from homologous protein clusters (HPCs) generated at 99% identity and 100% coverage using MMseqs2 15.6f452 [29]. PTU-specific networks were constructed with AcCNET [38], at 80% identity and 80% coverage. All networks were visualized in Gephi v0.9 with ForceAtlas2 layout.

### Pan-genome and enrichment analyses

PTU pangenomes were constructed with Roary v5.26.2 [39] with default parameters using GFF3 inputs. Gene–T6SS associations were tested with Scoary v1.6.16 [40], with FDR correction (p < 0.05). HPCs were annotated using eggNOG-mapper [41, 42] (e-value ≤ 1e-3, subject coverage ≥50%). Enrichment of COG categories in T6SS-positive plasmids vs. negative plasmids was tested by Fisher’s exact test (scipy.stats in Python v3.13), considering categories with FDR-corrected p < 0.05 and odds ratio >1 as enriched.

## Results

### Bacterial genomic landscape of T6SS distribution

To map the genomic distribution of T6SSs, we screened all bacterial chromosomes and plasmids from RefSeq212 using MacSyFinder, retaining genomes encoding ≥2 distinct T6SS components. Among 43,213 plasmids, 529 (1.2%) contained T6SS genes, largely within Proteobacteria, with sporadic occurrences in Bacteroidetes, Cyanobacteria, and Actinobacteria (Supplementary Table S5). Non-proteobacterial plasmids typically encoded partial clusters composed of *tssI/vgrG*, *tssE*, and occasionally *tssD/hcp* or *evpJ/PAAR*. Chromosomal T6SSs were likewise concentrated in Proteobacteria (10,945 genomes), followed by Bacteroidetes and a few additional phyla (Supplementary Table S6). Although this mirrors RefSeq taxonomic composition, it indicates that T6SSs are broadly distributed and not restricted to chromosomal loci in Proteobacteria.

We defined putatively complete T6SSs as clusters containing at least eight distinct core components, a threshold derived from the distribution shown in Figure 1. Applying this criterion, we identified 405 complete systems across 375 plasmids, with some plasmids carrying up to three T6SSs (Supplementary Table S7). Nearly all belonged to Proteobacteria (n=371), except for two cases in Bacteroidetes and single representatives in Acidobacteria and Gemmatimonadetes. Among Proteobacteria, subtype i T6SS-encoding plasmids (n=403) were distributed across 20 host families, predominantly *Burkholderiaceae* (n=138), *Rhizobiaceae* (n=75), *Enterobacteriaceae* (n=55), *Campylobacteraceae* (n=19), *Rhodobacteraceae* (n=14), *Vibrionaceae* (n=13), and *Yersiniaceae* (n=13). Two plasmid-encoded subtype iii T6SSs were identified in Bacteroidetes (NZ_CP027233.1 and NZ_CP095063.1). Chromosomes carried 14,789, 148, and 293 clusters of subtypes i, ii, and iii, respectively, some genomes encoding up to six T6SSs (Supplementary Table S8). A few genomes harbored more than one subtype: for example, *Francisella* (four genomes) and *Dongshaea* (one genome) carried both T6SS^i^ and T6SS^ii^.

**Figure 1:**
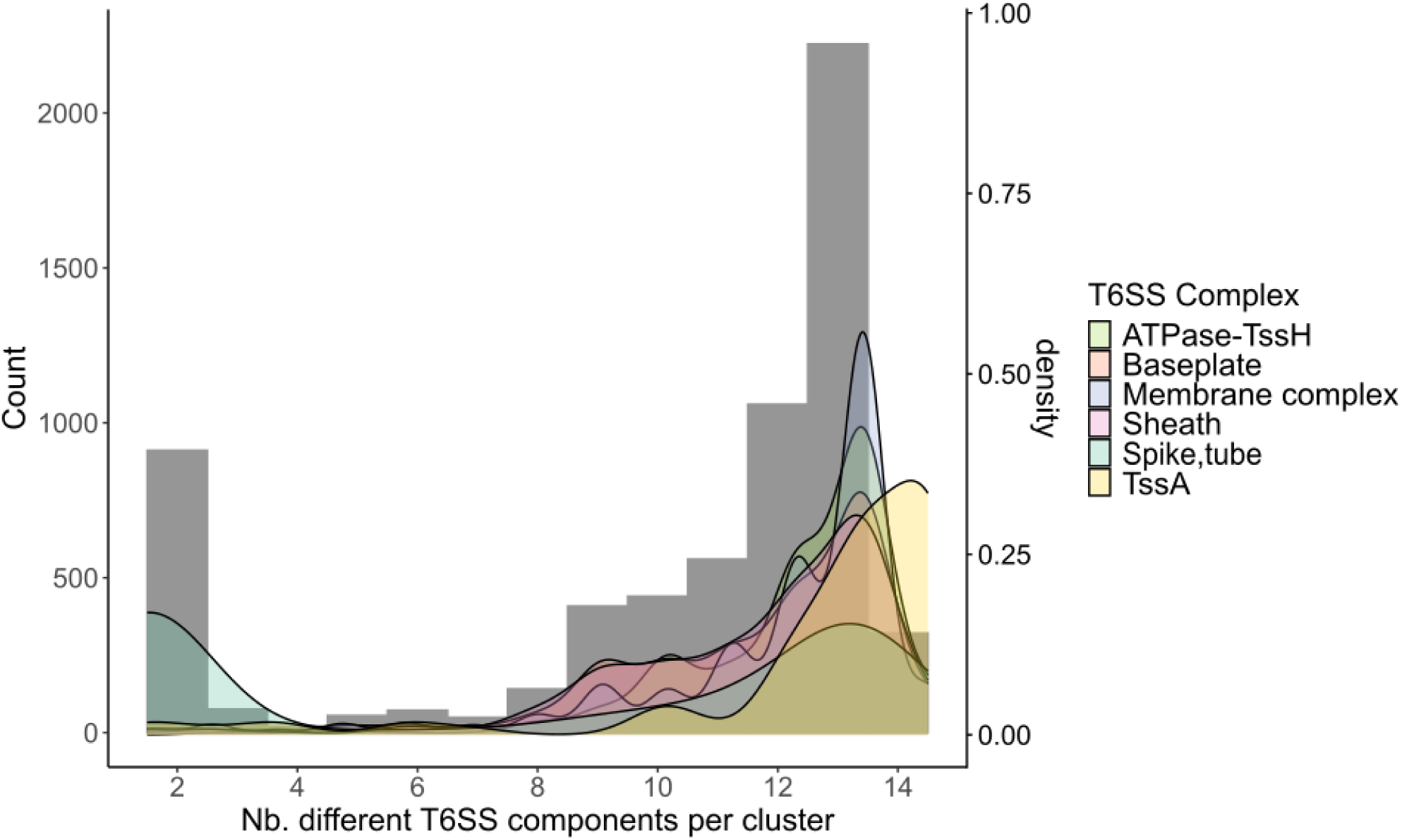
Composition of the detected plasmid-encoded T6SS^i^ clusters. The abundance of T6SS^i^ genes is shown according to the number of different components detected within the same cluster. Genes included in each T6SS subcomplex are as follows: *tssH* (ATPase); spike and tube (*evpJ/paar, tssI/vgrG,* and *tssD/hcp*); membrane complex (*tssJ, tssL, tssM*); baseplate (*tssE, tssF, tssG, tssK*), sheath (*tssB, tssC*); *tssA*.

In addition to complete systems, compact or truncated clusters were common. They typically encompass 2-3 genes combining *tssD/hcp* and spike components (*tssI/vgrG* and/or *evpJ/PAAR*) outside canonical T6SS clusters (Figure 1). We identified 467 such loci on 239 plasmids and 7,682 on 4,806 chromosomes. Of the 239 plasmids, 109 lacked a complete T6SS but carried one to three of these smaller loci. Proportionally, these reduced genomic arrays were more prevalent in plasmids than in chromosomes (54% vs. 34%), indicating greater modularization of T6SS elements in mobile replicons. These *hcp/vgrG/PAAR* loci, sometimes encoding only *hcp or vgrG*, correspond to orphan islands [43, 44]. These auxiliary modules are considered indicative of potential T6SS effector genes, as effector-encoding and secretion-related genes are frequently co-localized within these regions [45, 46]. To further assess this, we searched for isolated individual *hcp* and *vgrG* genes, identifying 299 plasmids (183 with *vgrG* and 116 with *hcp*). Among them, 163 lacked both complete T6SSs and larger orphan islands. Altogether, 420 plasmids carried orphan islands (*hcp, vgrG, hcp-hcp, vgrG-vgrG, hcp-vgrG, hcp-PAAR, vgrG-PAAR, hcp-vgrG-PAAR*), of which 272 encoded only these islands without an associated complete T6SS (Supplementary Table S7). These findings indicate that plasmids frequently mobilize individual *hcp* and *vgrG* modules independent of the full secretion apparatus, potentially facilitating modular exchange of effector genes.

### Plasmid genome size correlates with T6SS presence and complexity

Natural plasmids span a broad size range across bacterial phyla, often varying by three orders of magnitude [15]. Across all bacterial families, plasmids encoding complete T6SSs clustered toward the upper end of their size distributions and were consistently larger than T6SS-negative counterparts (Figure 2A). To contextualize this, we compared plasmid and host genome sizes. Using a ≥5% genome-size threshold to define megaplasmids [21] (Supplementary Table S3), 78.4% (294/375) of plasmids carrying complete T6SSs and 25.4% (69/272) plasmids with orphan islands met this criterion (Supplementary Table S7). Moreover, cumulative distribution analysis showed that plasmids encoding complete T6SSs (n=375) were significantly larger than those carrying only T6SS orphan islands (n=272; Mann-Whitney U test, *P*=2.63×10⁻^43^; Figure 2B).

**Figure 2:**
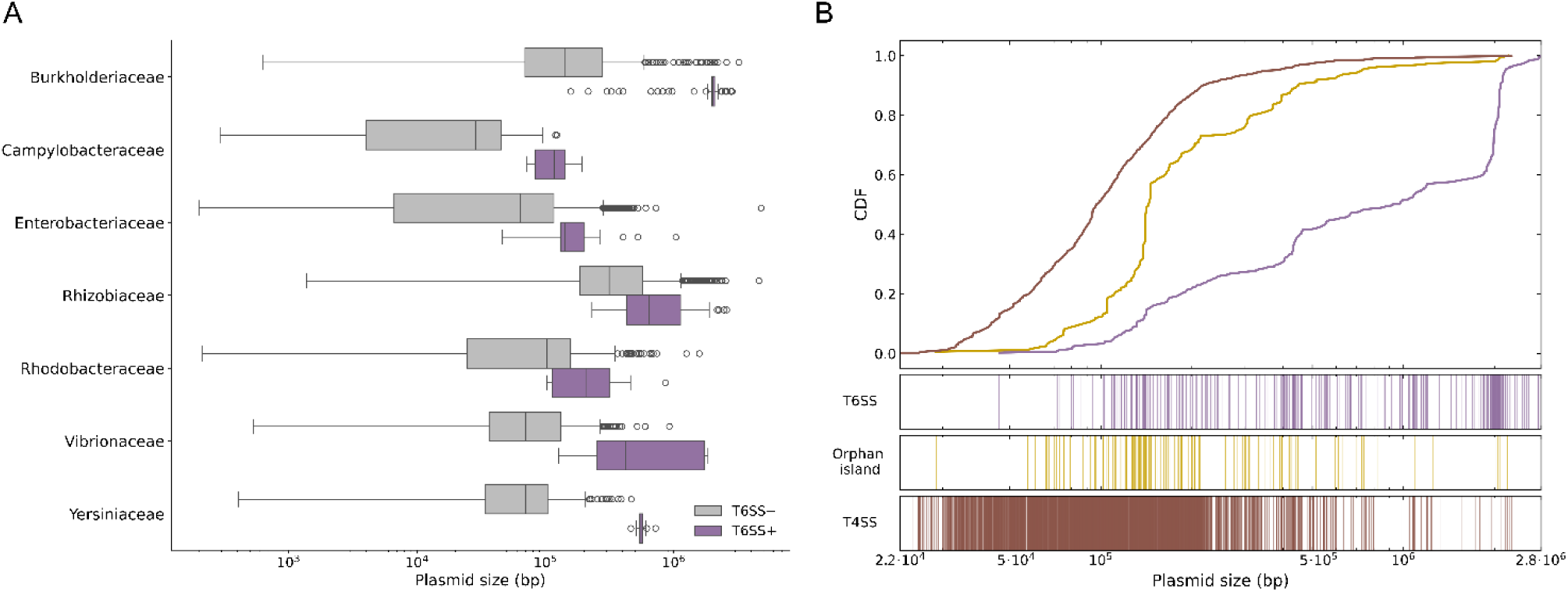
Genome size distribution of plasmids encoding T6SSs. A) Distribution of plasmid sizes for the most abundant bacterial families, differentiated by the presence (purple) or absence (gray) of a complete T6SS. Statistical differences were assessed using a one-sided Mann-Whitney U test for each family: *Burholderiaceae* (*P*=1.61×10⁻^61^), *Campylobacteraceae* (*P*=5.26×10⁻^13^), *Enterobacteriaceae* (*P*=1.22×10⁻^17^), *Rhizobiaceae* (*P*=1.15×10⁻^13^), *Rhodobacteraceae* (*P*=2.15×10⁻^5^), *Vibrionaceae* (*P*=3.75×10⁻^8^) and *Yersiniaceae* (*P*=3.52×10⁻^10^). B) Distribution of T6SSs, orphan islands, and T4SSs in bacterial plasmids. A total of 3,132 plasmids were ranked according to size (Supplementary Table S9). The top panel shows the CDF of genome size on a logarithmic scale for each plasmid subgroup. The bottom three panels display the presence of a complete T6SS (purple), orphan island (yellow) and T4SS (brown) for each genome. In all panels, the *x* axis represents size (bp) on log scale.

The T6SS operon typically comprises 13–14 core genes spanning >20 kb on average [47, 48], although some clusters are considerably larger [44]. While this substantial size could favor its retention on large plasmids, we tested whether operon length alone explains this association by comparing it to the type IV secretion system (T4SS), which contains a comparable number of genes (≈11 in the simplest MPF_T_ variant [49]). The cumulative size distribution of plasmids with T6SS was significantly shifted toward larger sizes relative to plasmids with T4SS (Mann-Whitney U test, *P*=1.14×10⁻^131^; Figure 2B), and only 6% of the latter group met the megaplasmid threshold (Supplementary Table S9), indicating that operon size alone cannot account for the strong link between T6SS presence and large plasmid size. These results suggest that additional functional or adaptive constraints, such as energy cost, stability, or selective advantage, may drive the preferential maintenance of T6SSs on large plasmids.

### Genomic characteristics of T6SS-encoding plasmids

To determine whether distinctive genomic features of large T6SS-encoding plasmids influence bacterial ecology and evolution, we analyzed plasmids carrying either complete T6SS clusters or orphan islands. We evaluated their potential for horizontal transfer by screening for conjugation-associated modules, their maintenance strategies via partition and toxin–antitoxin systems, and their accessory gene content, including AMR genes and virulence factors.

Plasmids were classified as transmissible (conjugative when both relaxase and mating-pair formation systems were present, or mobilizable when only relaxase was detected) or non-transmissible (lacking both). Plasmids carrying only orphan islands were highly enriched in transmissible categories (62.1%, 169/272; 6.3% mobilizable, 55.8% conjugative), whereas a smaller fraction of those with complete T6SS clusters were transmissible (30.4%, 114/375; 6.4% mobilizable, 24% conjugative) (Figure 3A; Supplementary Table S7). This suggests that horizontal transfer is more advantageous for disseminating effector modules, while complete T6SSs are typically maintained stably within hosts.

**Figure 3:**
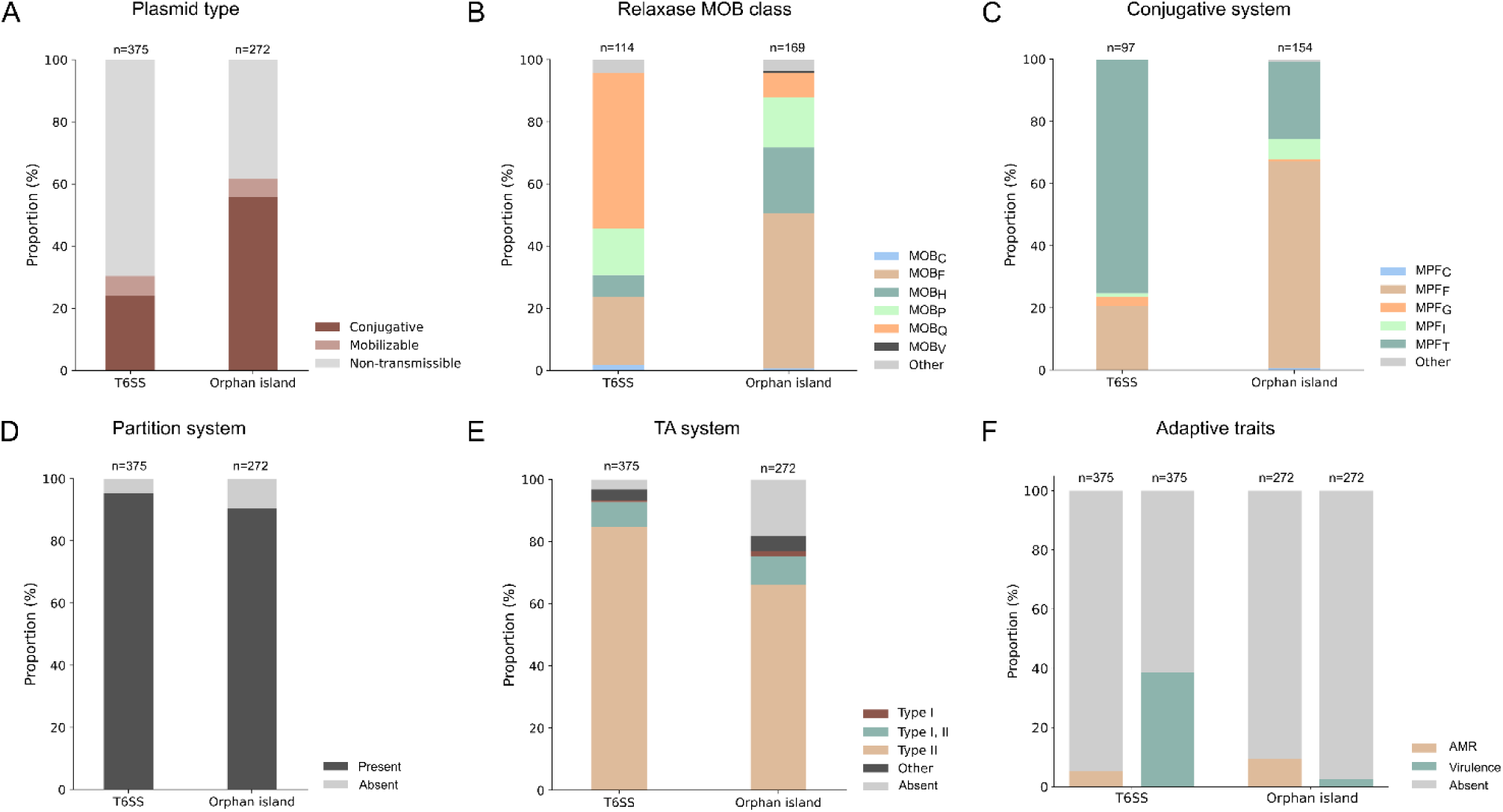
Mobility of plasmids encoding T6SSs. A) Proportion of conjugative, mobilizable and non-transmissible plasmids encoding a complete T6SS (n=375) or an orphan island (n=272), colored according to the legend. Proportion of plasmids encoding a complete T6SS or an orphan island according to B) the MOB relaxase class, C) the MPF type, D) partition system, E) TA module, and F) AMR and virulence traits.

Relaxase and conjugation system distributions were also different for plasmids encoding T6SS or orphan islands. MOB_F_ relaxases were most frequent (n=109), especially among orphan island plasmids (n=84; Fisher’s test, *P*=2.06×10⁻^6^), followed by MOB_Q_ (n=70), mainly linked to complete T6SSs (n=57; Fisher’s test, *P*=2.06×10⁻^6^) (Figure 3B; Supplementary Table S7). MOB_P_ (n=45) and MOB_H_ (n=43) occurred less often, the latter mainly associated with plasmids encoding orphan islands (n=35; Fisher’s test, *P*=0.002). Among mating-pair formation systems, MPF_F_ and MPF_T_ were predominant (122 and 112 plasmids, respectively): orphan-island plasmids were mainly MPF_F_-associated (n=102; Fisher’s test, *P*=9.85×10⁻^13^), whereas complete T6SS plasmids preferentially carried MPF_T_ (n=73; Fisher’s test, *P*=8.04×10⁻^15^) (Figure 3C; Supplementary Table S7). These associations suggest that distinct conjugation machineries favor the mobilization of different T6SS cargo types, shaping their evolutionary trajectories.

Partition systems were widespread, supporting stable inheritance of T6SS plasmids regardless of mobility potential. Most plasmids encoding complete clusters (358/375) and orphan islands (245/272) carried at least one partition protein, predominantly type I (ParA–ParB-like) (Figure 3D, Supplementary Table S7). TA modules, especially type II, were also widespread both in plasmids carrying complete T6SS (363/375, median=7) and orphan islands (223/272, median=2) (Figure 3E, Supplementary Table S7).

No significant association was observed between the presence of T6SS and AMR genes in plasmids. Only 45 encoded AMR determinants: 20 carrying complete T6SS and 25 with orphan islands (Figure 3F, Supplementary Table S7). Although most T6SS-encoding plasmids lacked AMR genes, those with orphan islands contained significantly more AMR determinants (median=8) than plasmids encoding complete T6SS clusters (median=1) (Mann-Whitney U test, *P*=2.57×10^−9^) (Figure 3F, Supplementary Table S7). Beyond AMR, we next examined the association between T6SS-encoding plasmids and virulence factors. These were more commonly associated with plasmids harboring complete T6SS clusters (145 of 375 plasmids) than with plasmids harboring orphan islands (7 of 272 plasmids; Fisher’s test, *P*=1.28×10⁻^29^) (Figure 3F). The detected virulence genes were primarily associated with exotoxins and adherence mechanisms: *flgG* (n=89), *fliI* (n=87), *flgI* (n=77), *cheW* (n=30), *fliP* (n=28) and *cheR* (n=25) (Supplementary Table S7).

The ANI network of 375 T6SS-positive plasmids (Supplementary Figure S1) revealed both densely connected groups and numerous singletons, reflecting high diversity and lineage-specific enrichment. Among these, 198 plasmids grouped into 13 PTUs across Proteobacteria (Supplementary Table S7), with 10 PTUs showing host range grade II (Supplementary Figure S2), suggesting that most T6SS-encoding PTUs can circulate among different species within the same genus. Each PTU exhibited a characteristic syntenic organization of its T6SS gene cluster (Supplementary Figure S3). In contrast, 84 of the 272 plasmids carrying orphan islands were assigned to 26 PTUs across Proteobacteria, encompassing host range grades I–IV (Supplementary Figure S2), which points to a broader taxonomic distribution of orphan islands—extending up to families within a given order—than that of complete T6SS clusters.

T6SS-positive plasmids were enriched in metabolic functions (COG G, I, E), often intersecting with chromosomal pathways, as well as in inorganic ion transport (P) and occasionally energy production (C) (Supplementary Figure S4), resembling megaplasmid profiles [50]. Analyses restricted to megaplasmids showed significant enrichments for signal transduction (T) in *Vibrionaceae* and *Rhodobacteraceae*, and amino acid metabolism (E) in *Rhizobiaceae*, suggesting selective coupling with T6SS carriage (Supplementary Figure S4).

### Ecological and evolutionary analysis of T6SS-encoding PTUs

Horizontal gene transfer via plasmids offers bacteria a rapid route to acquire competitive traits such as the T6SS, a contact-dependent nanoweapon that can shift competitive balances in microbial communities. However, the ecological significance of T6SS carriage at the level of persistent plasmid lineages (*i.e.* PTUs) remains underexplored. Here, we resolve relationships at the PTU level to assess whether specific plasmid lineages act as stable reservoirs, transient vectors, or evolutionary innovators of T6SSs.

We constructed PTU-level bipartite networks linking plasmids to homologous protein clusters (HPCs) (Supplementary Figure S5). This framework revealed whether T6SSs formed part of a PTU’s core proteome (≥80% of members), were sporadically present, or associated with distinct ecological subgroups. Several PTUs, including PTU-Bur3, PTU-E78, PTU-E79, and PTU-Rhi11, carried T6SSs as core features, indicating long-term stability. Others, such as PTU-Rhi3 and PTU-Rhi13, contained only a single T6SS-bearing member, consistent with sporadic acquisition, whereas PTU-Cam1 and PTU-Rhi4 exhibited intermediate patterns with T6SS-positive plasmids forming distinct subgroups, suggesting lineage-specific adaptation or niche specialization (Figure 4). The subgroup-specific segregation observed in PTU-Cam1 and PTU-Rhi4 makes them valuable models for understanding how plasmid-encoded T6SSs drive diversification. To explore these dynamics, we integrated PTU-specific core genome phylogenies with pan-genome-wide association analyses (pan-GWAS) to identify accessory traits linked to T6SS carriage.

**Figure 4:**
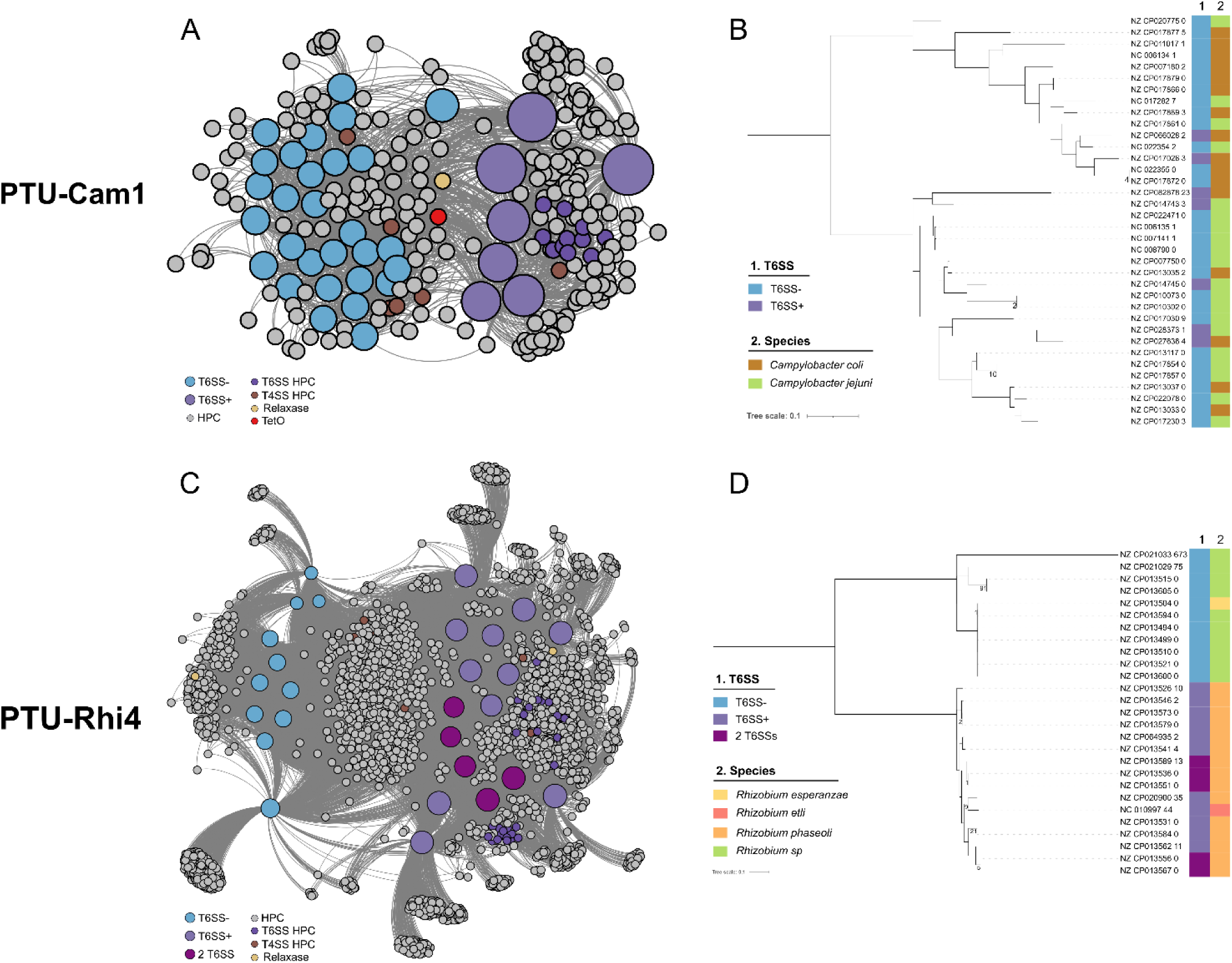
Features of PTU-Cam1 and PTU-Rhi4. A) Bipartite proteome network of PTU-Cam1 performed with AcCNET at 80% identity and 80% coverage. Large nodes represent plasmids, colored according to the presence of T6SS, while smaller nodes depict homologous protein clusters (HPCs). Most HPCs are colored in gray, while some corresponding to specific traits are colored according to the legend. B) Core-genome phylogenetic tree of PTU-Cam1. Parsimony tree based on core SNPs. Branch length scale represents changes per number of SNPs. Numbers at the internal nodes indicate the SNPs shared exclusively by their descendant lineages. The tree was midpoint-rooted. Bars to the right of the tree indicate (1) T6SS presence and (2) host species. C) Bipartite proteome network of PTU-Rhi4, obtained and visualized as described for PTU-Cam1. D) Core-genome phylogenetic tree of PTU-Rhi4, inferred and displayed as described for PTU-Cam1.

PTU-Cam1 represents a genus-restricted lineage of 36 *Campylobacter* plasmids (16 *C. coli*, 20 *C. jejuni*), of which seven encode complete T6SSs (4 *C. coli*, 3 *C. jejuni*). Network analysis (Figure 4A) revealed a clear division between T6SS-positive and -negative subgroups, the former clustering cohesively through shared HPCs. T6SS-positive plasmids were significantly larger (median 138 kb) than T6SS-negative members (median 46 kb) (Mann-Whitney U test, 2.7×10^−5^), qualifying as megaplasmids (≥5% of the *Campylobacteraceae* genome size). IS*200*/IS*605* transposases, found exclusively near T6SS loci, may mediate occasional acquisition events. Two conjugative megaplasmids previously shown to mediate hemolytic activity via T6SS (NZ_CP014745, NZ_CP014743; [10]) belong to this PTU, suggesting functional systems. The absence of phylogenetic separation between T6SS-positive and T6SS-negative members (Figure 4B) indicates that T6SS presence is not a primary driver of diversification. Nonetheless, pan-GWAS identified 99 genes associated with T6SS-positive plasmids, including uncharacterized loci, calcium/calmodulin-responsive adenylate cyclases, toxin–antitoxin systems, and phage integrases (Supplementary Table S10). PTU-Cam1 thus exemplifies a stable plasmid lineage in which the T6SS has become an integrated feature, reflecting co-adaptation with *Campylobacter* hosts and likely reinforcing intraspecific competition and influencing pathogenic interactions.

PTU-Rhi4 comprises 27 *Rhizobium* plasmids (15 *R. phaseoli*, 10 *Rhizobium* sp., one each *R. esperanzae* and *R. etli*), 16 of which carry a T6SS; five encode two distinct systems. In this PTU, T6SS carriage correlates with ecological divergence within the genus *Rhizobium*, as T6SS-positive plasmids are restricted to *R. etli* and *R. phaseoli*, whereas T6SS-negative plasmids occur in other *Rhizobium* species. As in PTU-Cam1, T6SSs are not part of the core proteome (Figure 4C), but T6SS-positive plasmids cluster together in the core-genome phylogeny, forming a distinct clade (Figure 4D), reflecting intra-PTU diversification structured at the intra-genus level. The two T6SS variants differ in distribution: one is shared with PTU-Rhi3 and several unassigned *Rhizobium* plasmids, while the other is shared with PTU-Rhi6 and PTU-Rhi13; both variants are confined to *Rhizobium*, suggesting variant-specific ecological roles. T6SS-positive plasmids are significantly larger (median 1.1 Mb) than T6SS-negative ones (median 751 kb) (Mann-Whitney U test, *P*=7.76×10^−6^) suggesting expansion of accessory content. Pan-GWAS identified enrichment for genes involved in inorganic ion transport/metabolism (COG P) and intracellular trafficking/secretion (COG U) (Supplementary Table S11), consistent with roles in competition or symbiosis in the plant rhizosphere. Taken together, these observations indicate that T6SS acquisition in PTU-Rhi4 is associated with ecological differentiation among *Rhizobium* species, likely reflecting adaptation to micro-niches shaped by resource partitioning.

### Phylogenetic relationships between chromosomal and plasmid-encoded T6SSs

To investigate the diversity and evolutionary trajectory of plasmid-encoded T6SSs, we reconstructed a maximum-likelihood phylogeny of all non-redundant TssC subunits from clusters containing at least ten components. The resulting tree (Figure 5A) recovered the expected deep clades (subtype T6SS^iv^ (brown, *A. asiaticus* 5a2), Bacteroidetes-specific T6SS^iii^ (blue)) and the *Francisella*-associated T6SS^ii^ (green) nested within the broader T6SS^i^ lineage (yellow), confirming their evolutionary relationships. This analysis also revealed coexistence of subtype i and ii systems within single genomes of *Francisella* and *Dongshaea*. Nearly all plasmid-encoded TssC subunits belonged to subtype i, with only two clustering within subtype iii.

**Figure 5:**
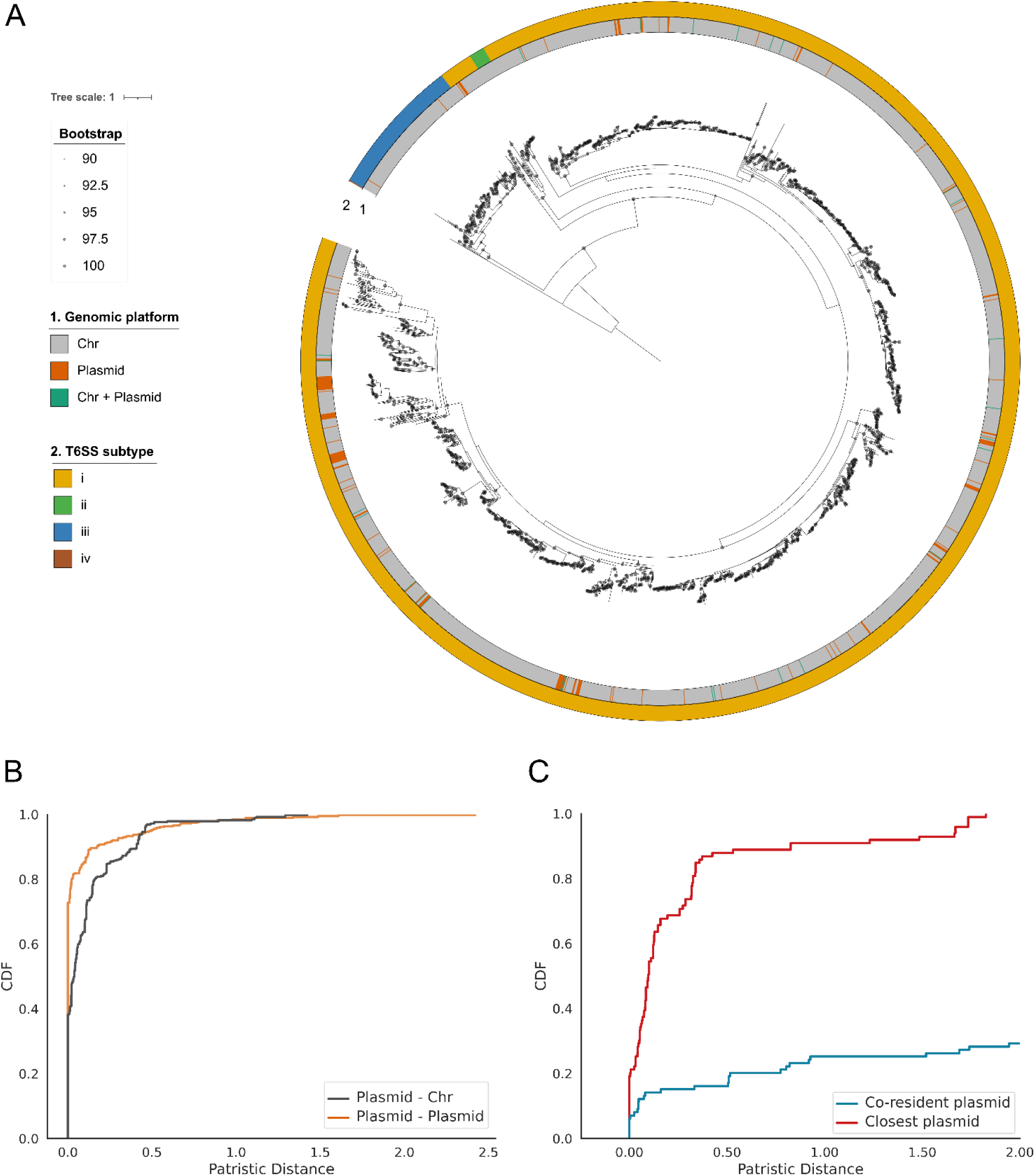
Phylogenetic analysis of T6SS. A) Maximum-likelihood tree of the TssC homologs retrieved from T6SSs comprising ≥10 different components. The tree is rooted with the TssC homolog gp18, the T4 phage sheath subunit. Nodes with UF-Bootstrap support values >=90% are indicated by gray circles. Rings, from inside to outside, indicate: 1) replicon (plasmid or chromosome) and 2) T6SS subtype. B-C) Cumulative distribution function of the patristic distances in the TssC phylogenetic tree. The curves represent the cumulative sum of the branch lengths linking each TssC to its closest homolog in the tree. B) For each plasmid-encoded TssC subunit, the distance to its closest homolog in another plasmid (orange) and in a chromosome (gray) is shown. C). For bacterial hosts encoding T6SS in both chromosome and plasmid (n=62), the minimal patristic distance from the chromosomal TssC to its co-resident plasmid homolog (blue) and to its closest plasmid homolog (red). Statistical differences were evaluated using a one-sided Mann-Whitney U test (p<0.001).

Plasmid-derived TssCs were scattered throughout the phylogeny, frequently interspersed with chromosomal homologs (inner ring, Figure 5A). Ancestral state reconstruction indicated recurrent plasmid acquisition from chromosomal sources (Supplementary Figure S6), followed by redistribution across diverse bacterial hosts (outer ring, Figure 5A), underscoring plasmids as major vectors of T6SS dissemination. Near-identical TssC proteins (≥99% identity, 100% coverage) occurred on both plasmids and chromosomes, providing clear evidence of recent horizontal transfer. The analysis also revealed multiple independent reintroductions of T6SSs from plasmids back into chromosomes (Supplementary Figure S6), highlighting the bidirectional nature of these exchanges. Importantly, nearly identical copies often spanned different species, genera, and even families, demonstrating that plasmids can breach deep taxonomic barriers during T6SS transfer.

Patristic distance analysis further underscored these dynamics. The CDF of patristic distances (Figure 5B) showed a steep initial rise for both plasmid–plasmid and plasmid–chromosome comparisons, with the former dominating. Of 385 plasmid-encoded TssCs, 70.1% (n=270) had identical homologs on another plasmid, consistent with tight clustering. Over half of these (53%; n=143) also had near-identical chromosomal homologs, indicating frequent exchange between compartments. Only 3.6% (n=14) matched exclusively to chromosomes. Even among non-identical pairs, plasmid–chromosome distances were short (median 0.11), suggesting ongoing transfers.

We next examined 62 isolates carrying T6SSs on both chromosomes and plasmids. For each, we compared the patristic distance between co-resident plasmid and chromosomal homologs to that between chromosomal TssCs and their closest plasmid homologs globally (Figure 5C). Co-resident pairs were rarely the closest relatives (Mann-Whitney U test, *P*=8.43×10^−20^), indicating dual systems arise from independent acquisitions rather than local duplications. These co-occurring systems likely broaden functional repertoires rather than increasing gene dosage, mirroring diversification among chromosomal T6SS loci.

A notable feature of these co-resident T6SSs was the high mobility of their plasmids: one-third (22/62) were conjugative, providing a direct route for cross-lineage transfer. Such plasmids likely mediate short-term ecological innovation by introducing new antagonistic systems while promoting long-term evolutionary stabilization. Supporting this, GC content of T6SS-encoding plasmids strongly correlated with that of their host chromosomes (Pearson’s correlation coefficient: r=0.94, *P*=1.44×10⁻³³; Supplementary Figure S7), with plasmids averaging ∼2% lower. This suggests that while plasmids act as mobile couriers of T6SSs, some achieve stable co-adaptation with their hosts over extended evolutionary timescales.

### Mechanisms of T6SS recruitment by plasmids

Phylogenetic analyses revealed extensive horizontal exchange of T6SSs between chromosomes and plasmids, with plasmid–plasmid transfers particularly frequent (Figure 5A; Supplementary Figure S6). To identify potential mechanisms, we examined the genetic context of T6SS loci for MGEs capable of mediating excision, mobilization, or integration, including phages, transposons, and insertion sequences (ISs). Specifically, we screened a 20-gene window around each chromosomal and plasmid-encoded T6SS for integrases, recombinases, and transposases.

Plasmids showed a marked enrichment of MGE signatures near T6SSs. Among 910 orphan islands and 403 complete T6SS^i^ clusters, over half contained at least one adjacent MGE (Supplementary Figure S8A). IS*5* and IS*3* were the most common, associated with both complete T6SSs and orphan islands (Fisher’s test, *P*>0.05), whereas IS*110* was largely restricted to orphan islands (Fisher’s test, *P*=9.13×10⁻^10^) (Supplementary Figure S8B). Chromosomal loci also harbored MGEs, but at lower frequencies: 3,347 of 14,789 complete T6SS^i^ and 2,863 of 7,209 orphan islands contained nearby ISs (Supplementary Figure S8A). On chromosomes, IS*3* (Fisher’s test, *P*=1.75×10⁻^5^) and IS*As1* (Fisher’s test, *P*=1.23×10⁻^30^) were primarily linked to orphan islands, while IS*5* occurred more often near complete clusters (Fisher’s test, *P*=3.41×10⁻^28^) (Supplementary Figure S8C).

These contrasting patterns underscore a major role for IS families, particularly IS*5* and IS*3*, in the mobility of T6SS loci. Their enrichment and close proximity to complete systems on plasmids suggest that these ISs act as molecular “handles” enabling plasmids to capture and redistribute T6SS clusters. Conversely, lower chromosomal frequencies imply that chromosomal loci function as reservoirs from which plasmids repeatedly acquire T6SS modules. Together, these results provide a mechanistic explanation for the widespread plasmid-mediated transfers inferred from phylogenetic analyses, identifying MGEs as key drivers of T6SS flux between chromosomes and plasmids.

### Recent transfer events between chromosomes and plasmids

To explore recent chromosomal–plasmid exchanges, we built bipartite networks linking plasmid- and chromosome-encoded T6SS^i^ systems containing ≥10 distinct components. The dataset comprised 342 plasmid-encoded and 12,792 chromosomal systems. Proteins were clustered at 99% identity and 100% coverage to identify near-identical homologs indicative of recent horizontal transfer. In the resulting network (Figure 6A), each T6SS^i^ was connected to all systems sharing protein clusters, such that highly interconnected communities reflect potential recent transfer events.

**Figure 6:**
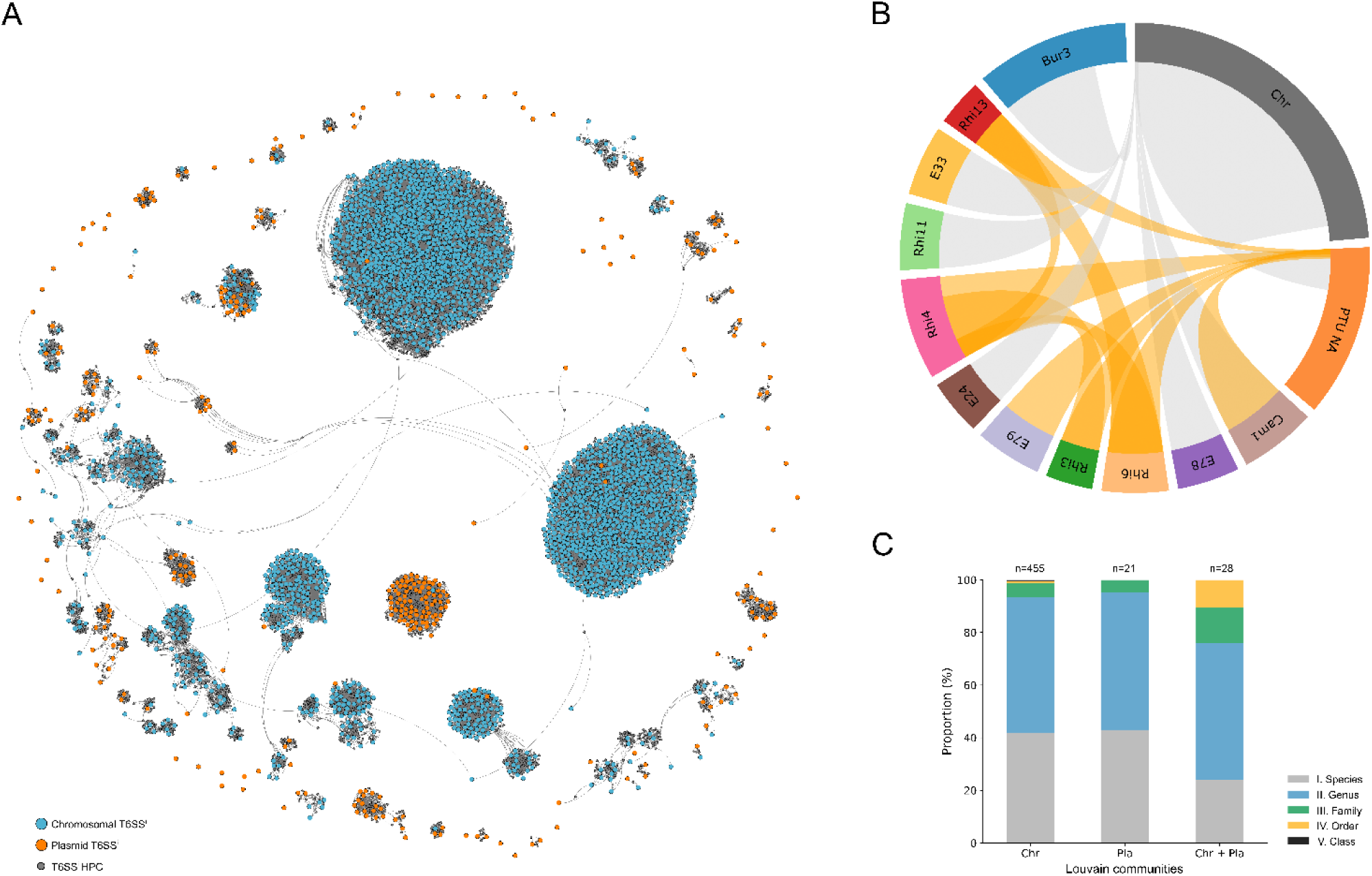
Recent horizontal gene transfer of T6SSs in plasmids and chromosomes. A) Bipartite network of plasmid and chromosomal T6SSs^i^ generated at 99% identity and 100% coverage. Nodes representing T6SS proteins are shown as small gray circles, while nodes representing complete T6SSs are larger and color-coded by replicon: plasmid (orange) and chromosome (blue). Only Louvain communities composed of plasmid T6SSs or mixtures of plasmid and chromosomal T6SSs are shown. B) Chord diagram showing connections in the network among PTUs, other plasmids (PTU NA), and chromosomes (Chr). Two genomic entities are linked by an edge when they belong to the same Louvain community. C) Host range of shared T6SS clusters detected within Louvain communities.

Community detection yielded 1,256 clusters (Supplementary Table S12). Over half (n=752) were singletons, including 57 plasmid-encoded systems—mostly from plasmids outside defined PTUs. Among the 504 multi-member communities, most comprised closely related systems, with near-zero patristic distances between TssC subunits (Supplementary Figure S9). Twenty-one communities contained only plasmid-encoded systems (n=129), and several joined plasmids from distinct PTUs (Figure 6B). For example, PTU-Rhi4 plasmids often carried two T6SSs—one clustering with PTU-Rhi3 and another resembling PTU-Rhi6 or PTU-Rhi13—indicating inter-PTU recombination or capture from shared chromosomal sources.

A total of 156 plasmid-encoded T6SSs grouped with chromosomal counterparts (Supplementary Table S12), mainly within PTU-Bur3, PTU-Cam1, PTU-E33, PTU-Rhi11, PTU-E78, and PTU-E24 (Figure 6B). Although chromosomal systems generally dominated these mixed communities, some (e.g., PTU-Cam1, PTU-Rhi11, PTU-E24) contained comparable plasmid and chromosomal representation. Of the 28 Louvain communities that included T6SSs on both chromosomes and plasmids, nine contained instances where the same transposase was located near the system on both replicons—for example, mixed communities featuring T6SSs on PTU-E33 plasmids and chromosomes, as well as on PTU-E78 plasmids and chromosomes. Most transfers occurred between closely related taxa, typically within species or genera, irrespective of genomic context (Figure 6C). Nevertheless, a minority of mixed communities spanned distinct families, indicating that while T6SS mobility is generally constrained by phylogenetic relatedness, plasmids can occasionally bridge deeper evolutionary divergences.

## Discussion

Our comparative genomic analyses reveal that T6SSs are not confined to chromosomal loci but are instead frequently encoded on plasmids, expanding current views of how machineries that mediate interbacterial antagonism are organized and propagated across bacterial genomes. From an evolutionary-ecological perspective, understanding plasmid-T6SS associations is crucial for elucidating how microbial populations balance horizontal mobility with competitive stability. We found that plasmid-encoded T6SSs are widespread across proteobacterial families but display a marked bias toward megaplasmids, indicating that these replicons provide suitable contexts for maintaining complex machineries.

Beyond their size, megaplasmids are characterized by chromosome-like features that stabilize low-copy replicons. In *Enterobacteriaceae* large plasmids encode ParAB partition systems and FtsK-dependent dimer resolution modules that safeguard faithful inheritance [51]. These mechanisms ensure correct segregation and tether plasmids to the nucleoid core [52], the site of replication, transcription, and recombination. Scaling laws in plasmid biology show that larger plasmids exhibit lower copy numbers and converge toward chromosomal organization [53], carrying tRNAs [54, 55], transcription- and translation-related genes [56], and nucleoid-associated proteins such as H-NS [57, 58]. Consequently, the strong association of complete T6SSs with megaplasmids likely reflects the capacity of these replicons to integrate, maintain, and disseminate complex adaptive traits.

The ecological consequences of T6SS carriage are context dependent. Some studies with chromosomal T6SSs report little or no fitness cost under laboratory conditions [59, 60], others reveal strong counter-selection in host-like or symbiotic environments [61–63]. This suggests that the benefits of megaplasmid-encoded T6SS are conditional, providing advantages in competitive or specific environmental contexts. We found that plasmids carrying orphan islands were more frequently associated with conjugative systems than those encoding complete T6SS loci (Figure 3), indicating that modular, T6SS-related cargo is particularly amenable to horizontal transfer. Consistently, plasmids with orphan islands exhibited broader host ranges than those carrying full T6SSs (Supplementary Figure S2). These orphan islands therefore act as flexible genomic platforms enabling rapid turnover of toxic effectors, promoting functional diversification without requiring mobilization of the entire secretion apparatus.

Within PTUs, T6SS distribution is highly heterogeneous. Some plasmid lineages retain T6SSs as stable, core features, whereas others exhibit sporadic or lineage-restricted occurrences. This diversity suggests multiple evolutionary trajectories, from long-term integration to transient acquisition. Certain families appear to have domesticated T6SSs, while others behave as transient vectors. Thus, plasmids function both as long-term repositories and short-term carriers of antagonistic potential.

The mobilization of complete T6SS loci likely reflects physical and genetic constraints on horizontal gene transfer. Typical transfer tracts of 25–30 kb per transfer event [64, 65] suffice to capture a full T6SS locus, yet such events are favored in the context of large MGEs [66]. Our phylogenetic analyses (Figure 5A and Supplementary Figure S6) indicated repeated acquisition of plasmid-borne T6SSs from chromosomal origins. This observation broadens our understanding of T6SS biology, showing that plasmids offer a structural context that overcomes the size constraints that restrict the transfer of full T6SS loci through natural transformation, which generally favors the acquisition of only a few genes at a time [66, 67]. Nearly identical T6SSs detected across distinct plasmid lineages and chromosomes (Figure 6), often flanked by transposases, offer direct evidence of ongoing bidirectional exchange across species, genera, and even taxonomic families. These recurrent transfers highlight plasmids as active agents breaching taxonomic barriers and accelerating the redistribution of complex adaptive systems. Similar dynamics underlie the spread of toxin and antibiotic-resistance modules [68, 69] emphasizing that plasmids serve simultaneously as transient vehicles and enduring scaffolds for antagonistic innovations. The ecological implications of this are substantial: a subset of plasmids encoding either T6SS or orphan islands in our dataset are conjugative (Figure 3), providing a mechanism for the rapid introduction of antagonistic or virulence traits into novel hosts and reshape microbial interactions across environments, from structuring competitive dynamics in different microbiomes to accelerating the emergence of clinical pathogens with enhanced virulence repertoires.

Our results support a mechanistic model in which plasmids act as both transient vehicles and long-term reservoirs of T6SSs, shaping bacterial competitiveness and adaptation across ecological niches. Functionally, plasmid-encoded T6SSs contribute to diverse ecological outcomes. In *Vibrio crassostreae* and *Rhizobium*, plasmid-borne T6SSs enhance host colonization or mutualistic symbiosis [9, 12, 70]. In commensal *Neisseria cinerea*, a plasmid-encoded T6SS enables outcompetition of related pathogens [11], while in *Campylobacter*, megaplasmid T6SSs enable erythrocyte lysis, facilitating persistence in blood-rich environments [10]. These examples illustrate that plasmid-encoded T6SSs can be ecologically maintained as antagonistic, symbiotic, or pathogenic tools depending on environmental context. In contrast, the preferential linkage of compact orphan islands to conjugative plasmids suggests a modular dissemination strategy, spreading specific effectors without the full apparatus. This division of labor mirrors an evolutionary balance between stability and mobility: megaplasmids provide long-term innovation hubs, whereas smaller plasmids accelerate effector turnover.

Collectively, our findings support a mechanistic–ecological model in which plasmids act as both reservoirs and vectors of T6SSs, shaping bacterial competitiveness and adaptation across environments. The preferential association of complete T6SSs with large, stably maintained plasmids underscores that these systems are not passive cargo but integrated components of plasmid–host consortia. By coupling stability with mobility, plasmids enable the continuous recruitment, refinement, and redeployment of antagonistic modules, influencing microbial community structure and evolutionary trajectories. Future work should explore how these dynamics affect community resilience, cooperation–competition balance, and the long-term diversification of microbial lineages.

## Data availability

The authors confirm all supporting data and protocols have been provided either within the article, in supplementary data files, or through supporting data files available on GitHub (https://github.com/mdmqc/Plasmid_T6SS).

## Acknowledgments

This work was supported by the Spanish Ministry of Science and Innovation (Grant MCIN/AEI/10.13039/501100011033 PID2020-117923GB-I00 to MPG-B) and the Spanish Ministry of Universities (predoctoral contract FPU20/04579 to MMQ-C).

