## Supplementary Figures for "“Plasmids drive the dissemination and diversification of Type VI Secretion Systems”"

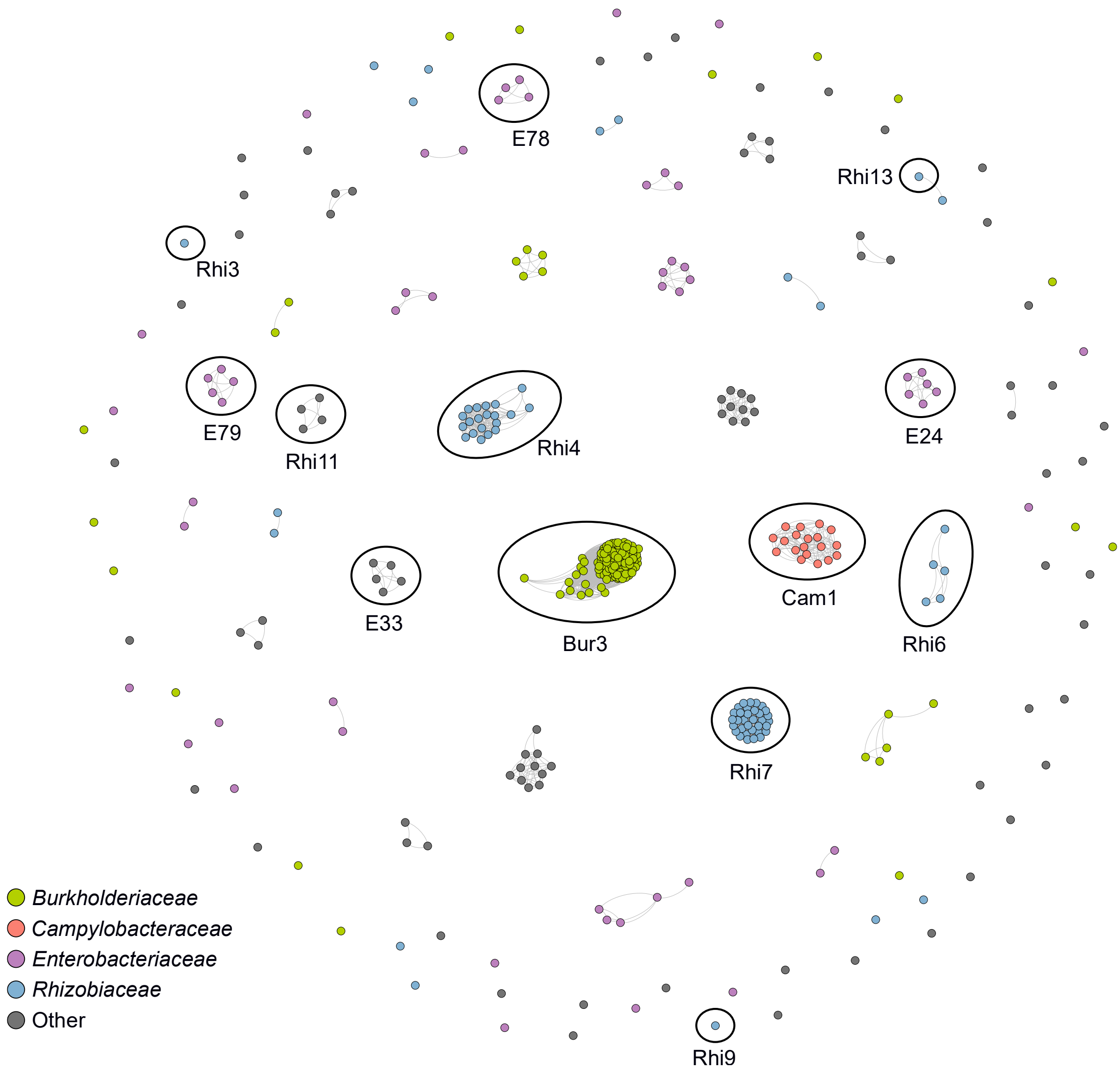


**Supplementary Figure S1. ANI$_{L50}$ similarity network of plasmids encoding complete T6SS.** Nodes, representing T6SS-encoding plasmids (n=375), are colored according to their host taxonomy, as indicated in the legend. Plasmid Taxonomic Units (PTUs) are encircled when assigned.

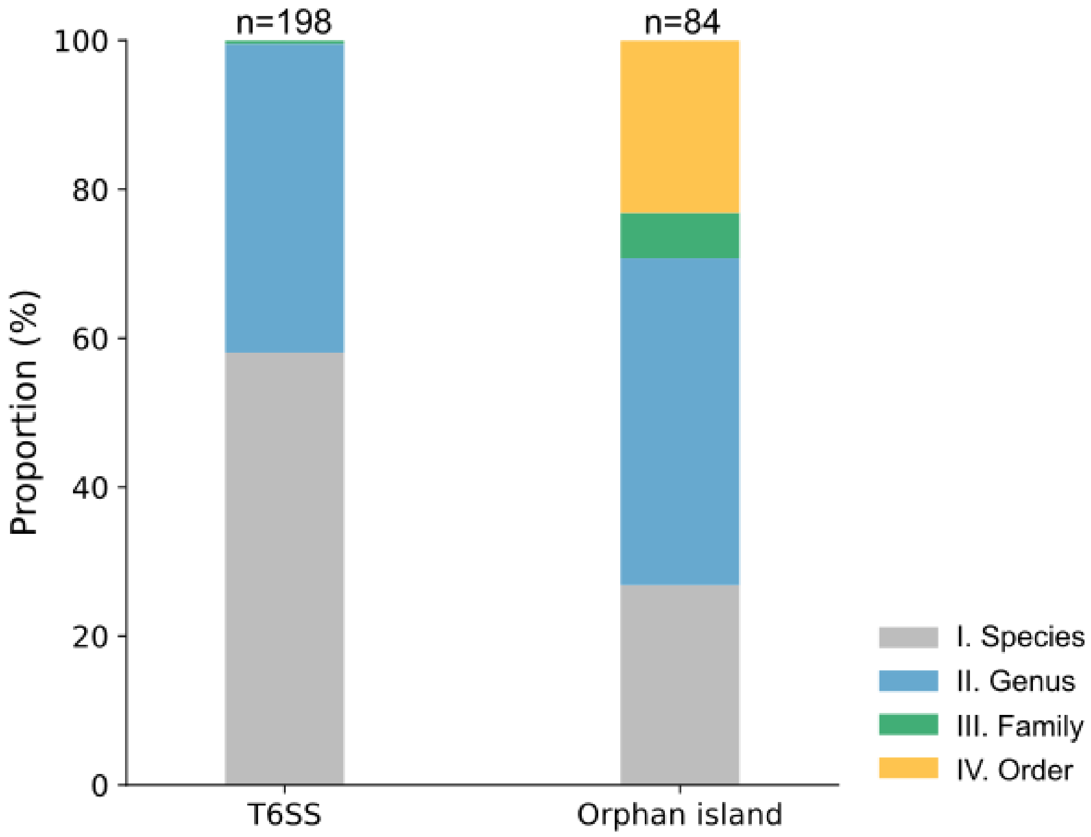


**Supplementary Figure S2. Host range of PTUs encoding T6SS.** The graph shows the taxonomic host distribution for PTUs encoding complete T6SSs (n=198) and orphan T6SS islands (n=84), colored according to the legend.

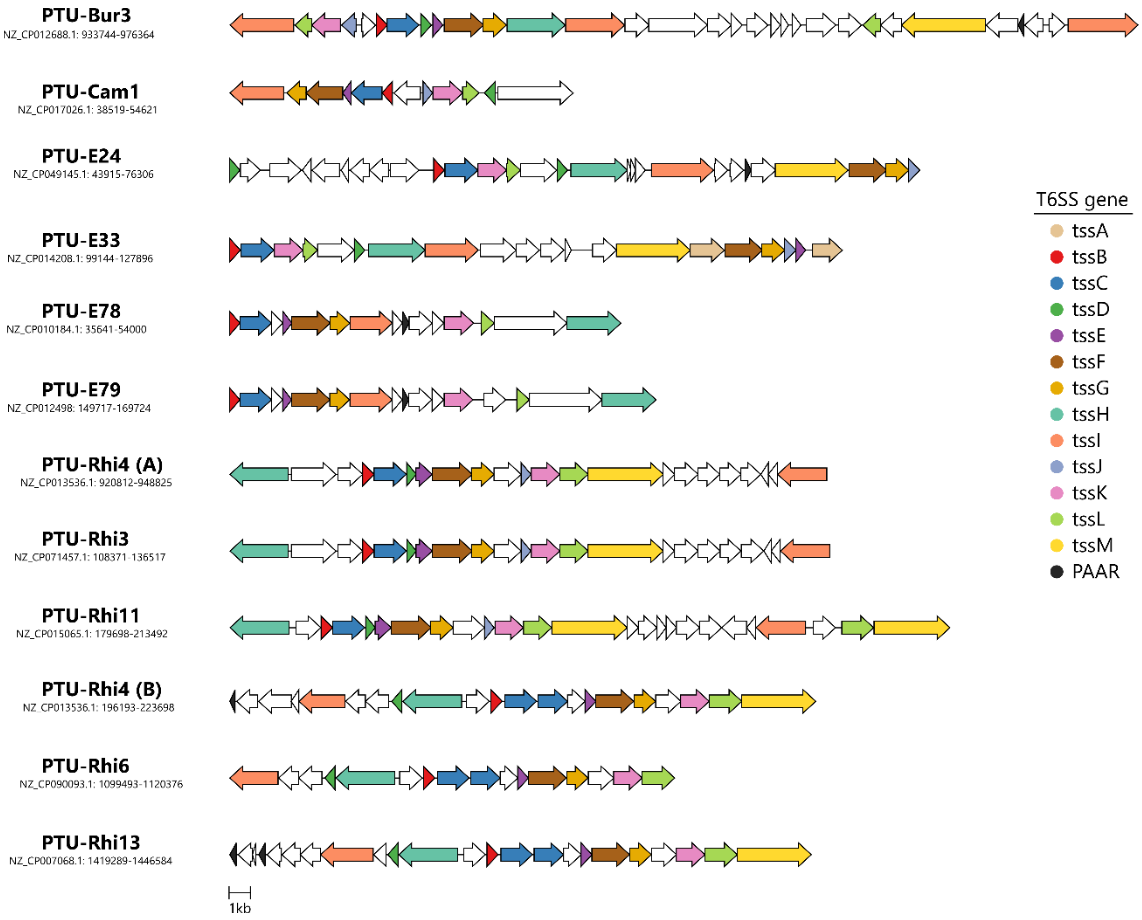


**Supplementary Figure S3. T6SS genetic organization in representative plasmids from different PTUs.** The synteny of the system is shown for each PTU encoding a complete T6SS using a representative genome (accession number and coordinates are indicated). T6SS genes retrieved from this study are colored according to the legend.

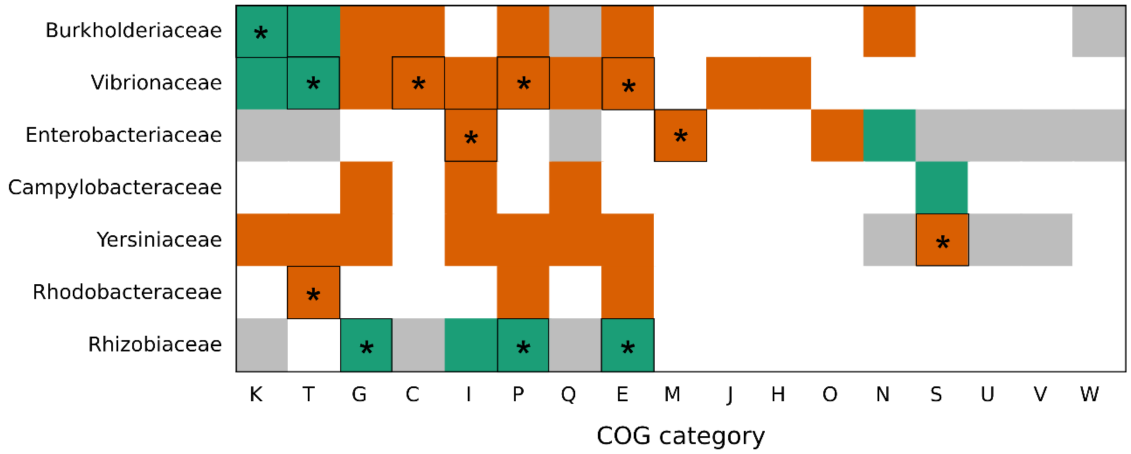


**Supplementary Figure S4. Functional COG categories enriched in genomes encoding complete T6SS.** For each host family, significantly enriched functional categories (Benjamini-Hochberg, p-adjust <0.05) are depicted in orange if exclusive to plasmids, gray if exclusive to chromosomes, or green if present in both replicons. COG categories significantly enriched preferentially in T6SS$^{+}$ megaplasmids relative to T6SS$^{-}$ counterparts are marked with (*). List of COG categories: C: energy production and conversion; E: amino acid transport and metabolism; H: coenzyme transport and metabolism; I: lipid transport and metabolism; G: carbohydrate transport and metabolism; J: translation, ribosomal structure and biogenesis; K: transcription; M: cell wall/membrane/envelope biogenesis; N: cell motility; O: post-translational modification, protein turnover, chaperones; P: inorganic ion transport and metabolism; Q: secondary metabolites biosynthesis, transport and catabolism; S: function unknown; T: signal transduction mechanism; U: intracellular trafficking, secretion and vesicular transport; V: defense mechanisms; W: extracellular structures.

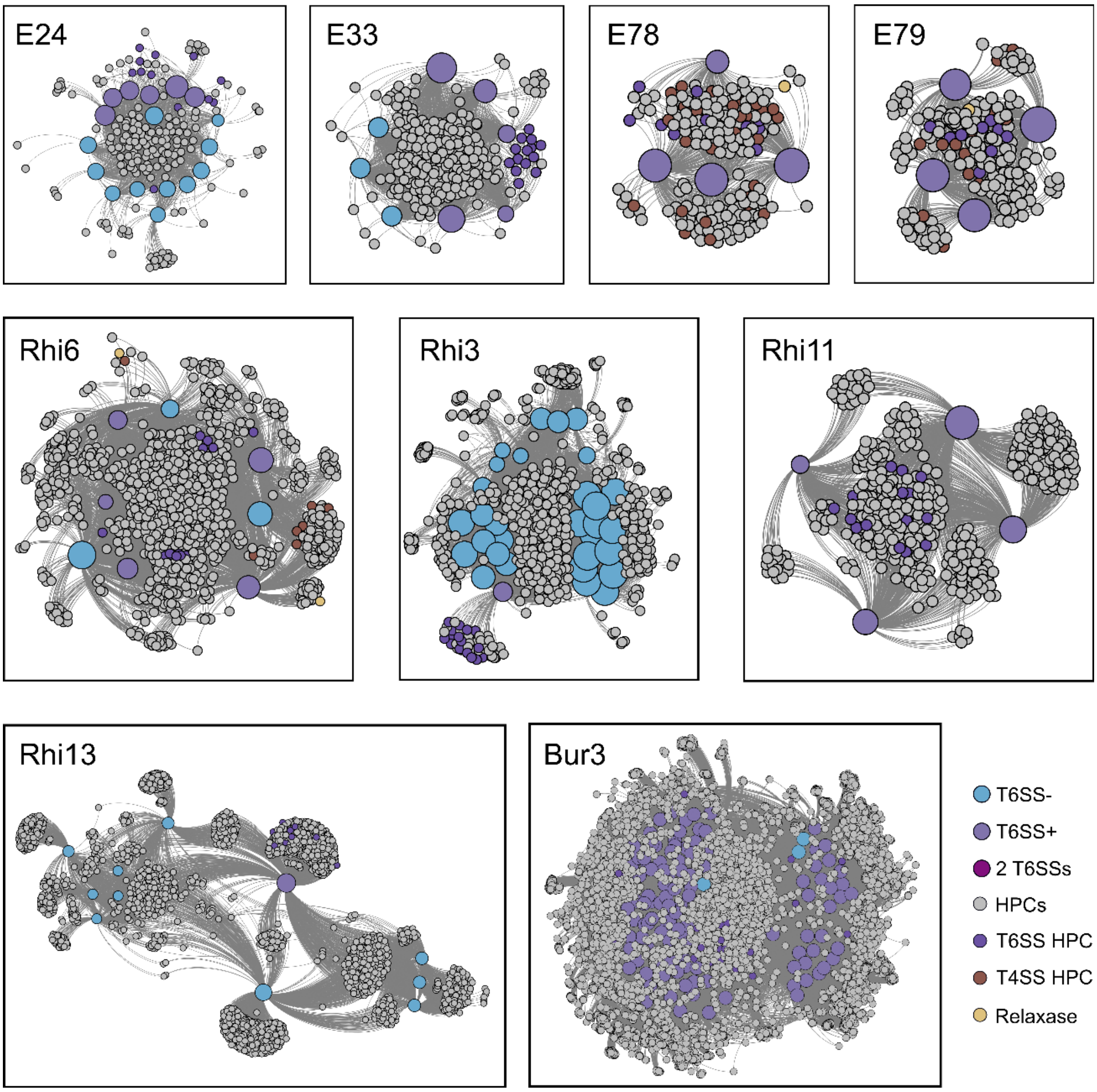


**Supplementary Figure S5. Proteome networks of the T6SS-encoding PTUs.** Bipartite proteome networks generated with AcCNET at 80% identity and 80% coverage for each PTU containing plasmids encoding complete T6SS in this study. Large nodes represent plasmids colored according to the presence of T6SS[i], while smaller gray nodes depict homologous protein clusters (HPC). HPCs corresponding to T6SS, T4SS, and relaxase components are colored according to the legend.

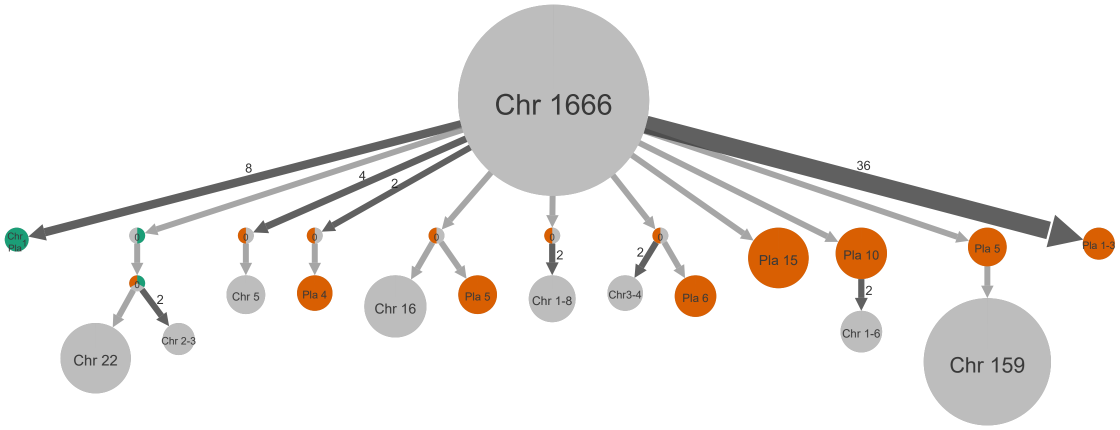


**Supplementary Figure S6. Inference of ancestral states of TssC subunits.** The ancestral states of TssC subunits from the phylogenetic tree (Figure 5) were inferred using PastML. The number of nodes contained in each state is shown, with chromosomal T6SS depicted in gray, plasmid-encoded TssC in orange and TssC found in both plasmid and chromosomal in green. Internal nodes shown as multi-colored circles containing “0” indicate ambiguous states for which no single ancestral state could be inferred. Numbers inside circles indicate the number of internal nodes or tree tips represented after vertical compression, whereas numbers on arrows indicate the number of identical subtrees merged horizontally. For a visual explanation of the interpretability of the ancestral compression, see the PastML help page (https://pastml.pasteur.fr/help).

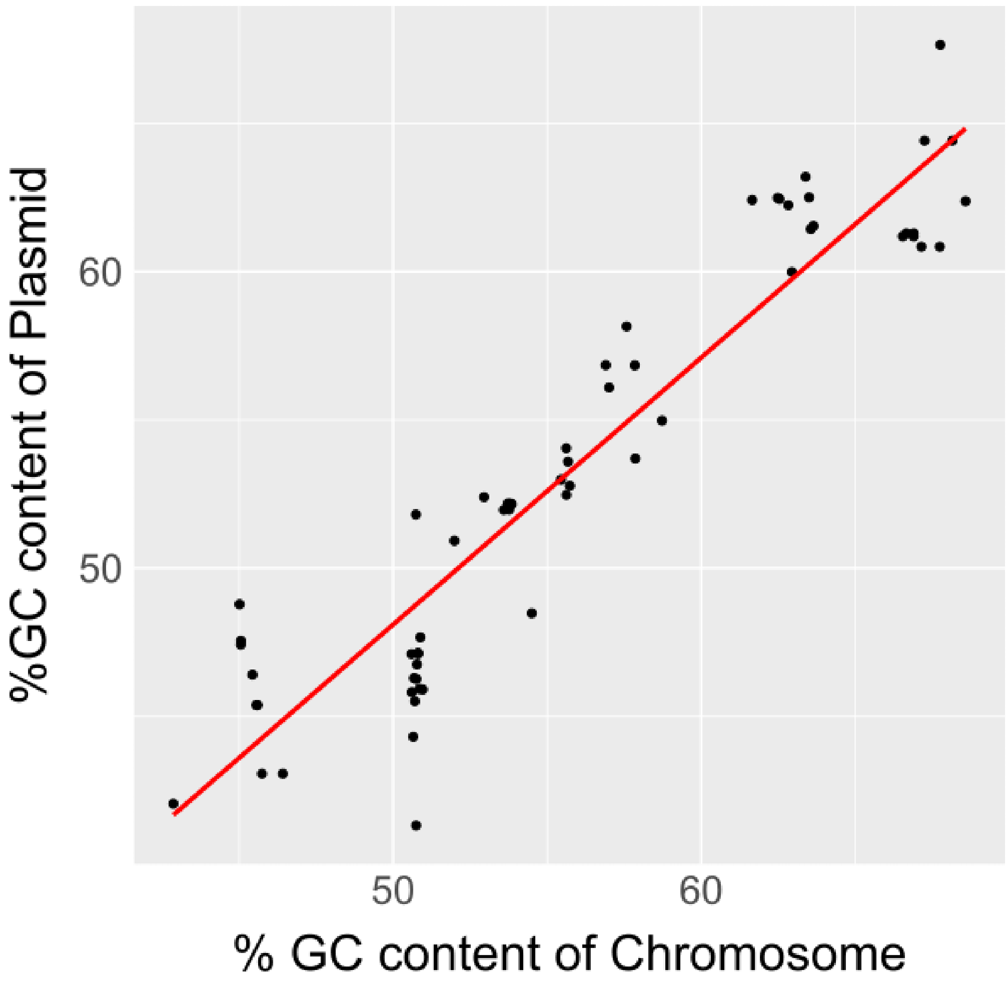


**Supplementary Figure S7. Comparison of GC content between co-resident plasmids and chromosomes encoding complete T6SSs.** %GC content of T6SS-encoding plasmids is plotted against %GC of their co-resident T6SS$^{+}$ chromosome (62 total comparisons). The red line indicates the linear regression model of the data (Pearson's correlation coefficient: r=0.94, p=$1.44\times10^{-33}$).

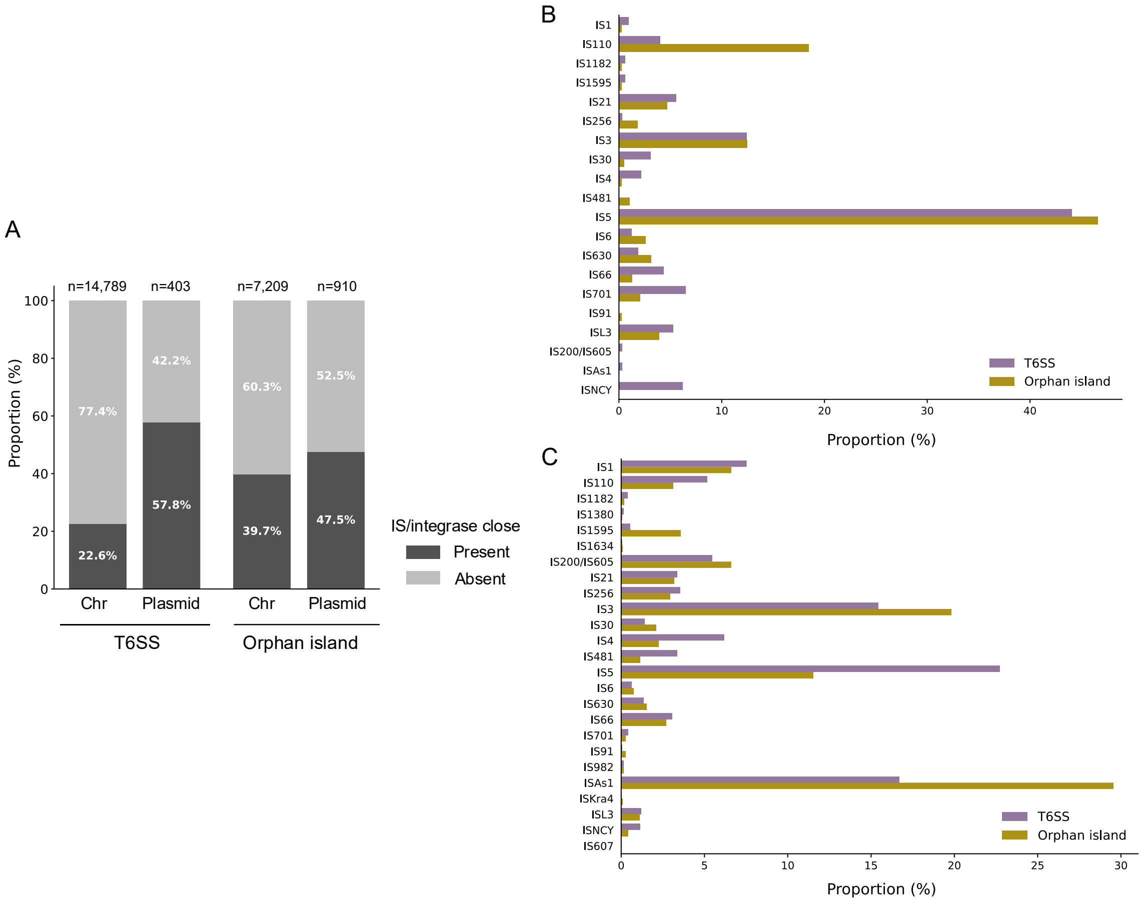


**Supplementary Figure S8. Presence of Insertion Sequences and integrases/recombinases in the proximity of T6SS[i].** A) Proportion of complete T6SS[i] clusters and orphan islands with an IS and/or integrase within 20 coding sequences from a T6SS component for chromosomes and plasmids. The number of T6SS[i] clusters analyzed in each case is indicated. B) Abundance of each IS family near a T6SS[i] or orphan island in plasmids. For each IS family, the proportion was calculated as the ratio of the number of ISs for that family in close proximity relative to the total number of ISs detected in close proximity to each cluster type (T6SS or orphan island). C) Abundance of each IS family near a T6SS[i] or orphan island in chromosomes. The proportion was calculated as in B.

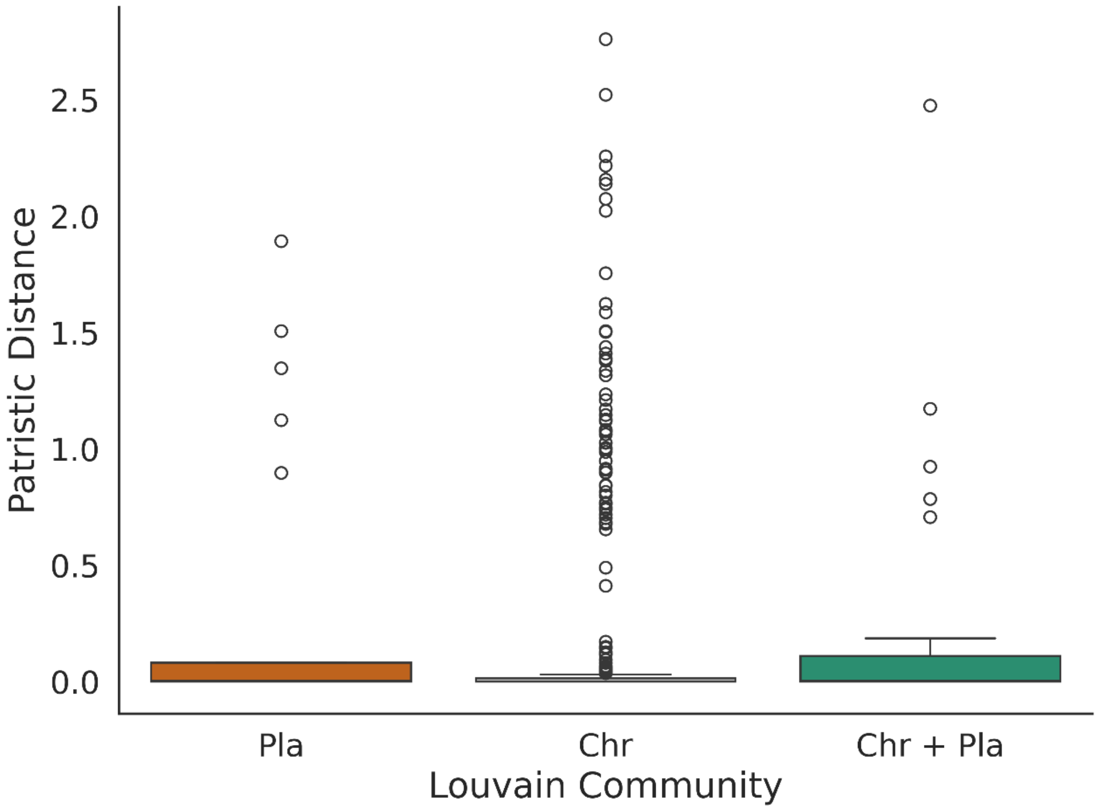


**Supplementary Figure S9. Phylogenetic relationships among T6SS$^{i}$ in the horizontal gene transfer network.** Distribution of patristic distances in the TssC tree (Figure 5A) for Louvain communities identified in the bipartite network (Figure 6A), considering T6SSs encoded exclusively on plasmids (Pla), exclusively on chromosomes (Chr), or on both plasmids and chromosomes (Chr + Pla).
